# Coccolith Sr/Ca is a Robust Temperature and Growth Rate Indicator that Withstands Dynamic Microbial Interactions

**DOI:** 10.1101/2021.10.28.466229

**Authors:** Or Eliason, Einat Segev

## Abstract

Coccolithophores are a diverse group of calcifying microalgae that have left a prominent fossil record on Earth. Various coccolithophore relics, both organic and inorganic, serve as proxies for reconstruction of past oceanic conditions.

*Emiliania huxleyi* is the most widely distributed representative of the coccolithophores in modern oceans, and is known to engage in dynamic interactions with bacteria. Algal-bacterial interactions influence various aspects of algal physiology and alter algal alkenone unsaturation (U^K’^_37_), a frequently used organic coccolithophore-derived paleotemperature proxy. Whether algal-bacterial interactions influence inorganic coccolithophore-derived paleo-proxies, is yet unknown.

A commonly used inorganic proxy for past productivity and sea surface temperature is the Sr/Ca ratio of the coccolith calcite. Interestingly, during interactions between bacteria and a population of calcifying algae, bacteria were shown to physically attach only to non-calcified algal cells, suggesting an influence on algal calcification.

In this study we explore the effects of algal-bacterial interactions on calcification and coccolith Sr/Ca ratios. We find that while bacteria attach only to non-calcified algal cells, coccolith cell coverage and overall calcite production in algal populations with and without bacteria, is similar. Furthermore, we find that Sr/Ca values are impacted only by water temperature and algal growth rate, regardless of bacterial influences on algal physiology. Our observations reinforce the robustness of coccolith Sr/Ca ratios as a paleo-proxy independent of microbial interactions, and highlight a fundamental difference between organic and inorganic paleo-proxies.

**Summary Statement:** The current research investigates the effect of microbial interactions on coccolith Sr/Ca ratio and overall calcification in the coccolithophore *Emiliania huxleyi*. We co-cultured *E. huxleyi* with the marine bacterium *Phaeobacter inhibens* and compared coccolith Sr/Ca between different growth stages in a range of temperatures. Our results indicate that coccolith Sr/Ca depends on temperature and algal growth rate, and remains robust despite significant bacterial influences on algal physiology.

## Introduction

Coccolithophores are a highly diverse group of single-celled marine algae belonging to the Haptophyte clade (Jordan and Chamberlain 1997, de Vargas, Aubry et al. 2007). They evolved over 200 million years ago, and have flourished in the world oceans ever since (Bown 1985, Mai, von Salis Perch-Nielsen et al. 1997). Their name is derived from the elaborate CaCO_3_ platelets, termed coccoliths, that cover the algal cell (Young, Didymus et al. 1992). The coccoliths are produced intracellularly in a designated compartment termed the coccolith vesicle, and are ejected onto the cell surface upon completion (Van Der Wal, De Jong et al. 1983, Marsh 2003, Gal, Wirth et al. 2016, Brownlee, Langer et al. 2020). Once exocytosed, the coccoliths adjoin one another to form a continuous shell.

Coccolithophore relics, both mineral and organic, are preserved in sediments and can provide chemical evidence of past environmental conditions (Stoll and Ziveri 2004, Baumann, Andruleit et al. 2005). Given the long evolutionary history and vast global distribution of the coccolithophore group, these primary producers have left behind one of the most extensive records of marine paleo-proxies, often dominating the bulk CaCO_3_ content in marine sediments (Broecker and Clark 2009, Bordiga, Cobianchi et al. 2014). An example for a commonly used haptophyte-derived organic paleo-proxy is the alkenone unsaturation index (U^K’^_37_), providing information about past sea surface temperatures (SST) (Brassell, Eglinton et al. 1986, Prahl and Wakeham 1987, Marlowe, Brassell et al. 1990).

A prevalent inorganic coccolithophore-derived paleo-proxy is the Sr/Ca ratio in the CaCO_3_ coccoliths, offering evidence for past productivity and SST. Coccolith Sr/Ca ratio has been studied both in sediments and culture experiments, revealing a connection between algal growth, calcification rates, and Sr/Ca values (Stoll and Schrag 2000, Stoll and Schrag 2001, Stoll, Klaas et al. 2002, Stoll, Rosenthal et al. 2002, Stoll, Ziveri et al. 2007, Müller, Lebrato et al. 2014, Mejía, Paytan et al. 2018). In laboratory experiments, a strong influence of temperature on Sr/Ca has been observed (Stoll, Klaas et al. 2002, Stoll, Rosenthal et al. 2002, Müller, Lebrato et al. 2014), similar to other biogenic CaCO_3_ sources (Rosenthal, Boyle et al. 1997, Goodkin, Hughen et al. 2005, Freitas, Clarke et al. 2006).

Accumulating studies indicate that coccolithophore physiology, and consequently coccolithophore remains, are largely influenced by biotic interactions, especially with bacteria (Harvey, Deering et al. 2016, Segev, Wyche et al. 2016, Barak-Gavish, Frada et al. 2018, Whalen, Kirby et al. 2018). To study these influences, we have previously established a model system for the co-cultivation of coccolithophores and bacteria. The algal-bacterial pair, consisting of the coccolithophore *Emiliania huxleyi* and the bacterium *Phaeobacter inhibens*, was chosen according to environmental data indicating that these specific algal and bacterial species co-occur in the marine environment and are likely to interact (Green, Echavarri-Bravo et al. 2015, Segev, Wyche et al. 2016).

Laboratory studies revealed that *E. huxleyi* and *P. inhibens* engage in a dynamic interaction, mediated through the exudation of various metabolites into the surrounding environment (Segev, Wyche et al. 2016). Initially the interaction is beneficial for both partners; algae exude dissolved organic matter (DOM) essential for bacterial growth. Consequently, bacteria secrete a hormone that stimulates algal growth, resulting in higher final densities of the algal population. As the algal population senesces, bacteria switch from being mutualistic to being pathogenic and kill their algal partners. The bacterial gain from killing the algal host is most likely the release of algal cell content, providing bacteria additional nutrients (Segev, Wyche et al. 2016, Barak-Gavish, Frada et al. 2018). Importantly, the bloom-and-bust dynamics observed in laboratory experiments are similar to natural *E. huxleyi* bloom dynamics in the ocean (Tyrrell and Merico 2004, Behrenfeld and Boss 2014).

In light of the bacterial impact on algal physiology and the environmental relevance of this microbial interaction, previous studies explored the influence of algal-bacterial interactions on algal relics that serve as paleo-proxies, specifically the alkenone unsaturation index. Culture experiments indicated that algal-bacterial interactions result in a measurable influence on U^K’^_37_, significantly modifying temperature reconstructions (Segev, Castaneda et al. 2016).

The algal-bacterial relationship involves a physical aspect; In previous reports, bacteria were observed to attach directly onto the algal cells (Segev, Tellez et al. 2015, Segev, Wyche et al. 2016, Barak-Gavish, Frada et al. 2018). Interestingly, when bacteria were cultivated with a calcifying *E. huxleyi* strain, in which the extent of coccolith coverage, i.e. the number of coccoliths covering each cell, varies within the population, physical attachment of bacteria onto algae was restricted to uncalcified cells exhibiting no coccoliths at all, termed naked cells. Algal cells covered by coccoliths exhibit no attached bacteria (Segev, Wyche et al. 2016). Despite the natural variability in coccolith coverage, these observations raised the possibility that *P. inhibens* bacteria might influence algal CaCO_3_ production or its retainment on the cell surface.

Here, we sought to explore the bacterial influence on coccolithophore CaCO_3_ production or cell coverage and derived inorganic remains, namely coccoliths Sr/Ca ratio. Culture experiments were conducted under different temperatures, in which marked changes in algal-bacterial dynamics were evident. Our findings reveal that bacteria do not influence coccoliths production or coverage. Furthermore, our results indicate that coccolith Sr/Ca depends on temperature and algal growth rate, regardless of significant bacterial influences on algal physiology.

## Results

To assess possible bacterial influences on the elemental composition of algal coccoliths, we first assessed the general bacterial impact on algal calcification. As previously reported, in algal-bacterial co-cultures, bacteria appear to be exclusively attached to naked algal cells (Segev, Wyche et al. 2016), and not to calcified algal cells that are present in the culture (Fig. 1a, b, c, d). To determine whether bacteria inhibit algal calcification, or whether bacteria attach to pre-existing naked algal cells, we evaluated if bacteria affect algal calcite production in co-cultures. As can be seen in Figure 1, when both axenic and co-cultures are subjected to centrifugation using a viscous solution that promotes coccolith separation from biomass and accumulation at the bottom of a tube (see sample preparation in Materials and Methods), all cultures exhibit a significant white pellet (Fig. 1e). Inspection under a scanning electron microscope (SEM) revealed that the white pellet in all cultures is composed of coccoliths (Fig. 1f). Thus, it appears that coccoliths are produced both in axenic and co-cultures, and no major bacterial influence on algal calcification is observed.

**Figure 1.**
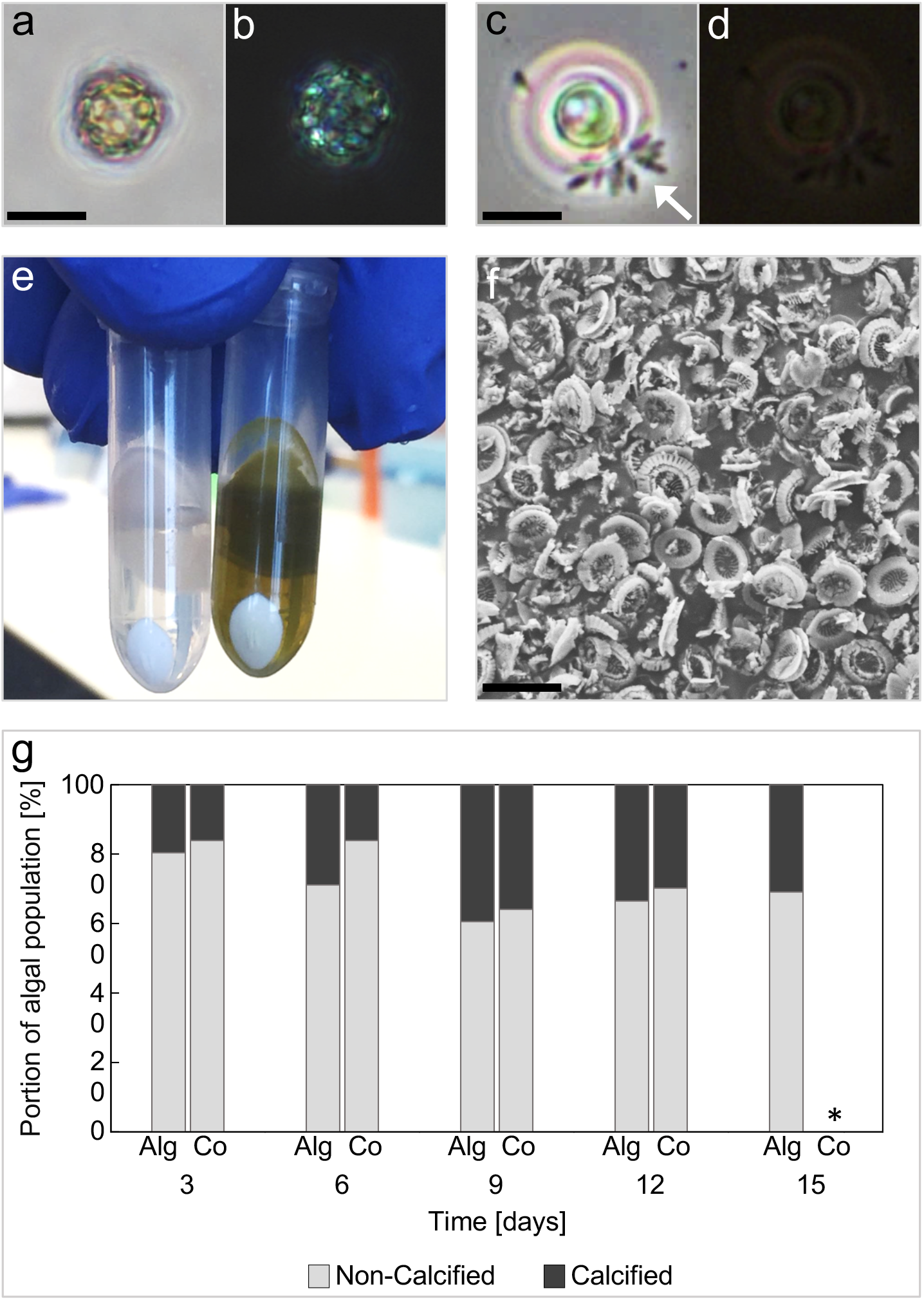
Calcification and coccolith coverage in *E. huxleyi* are not affected by microbial interactions. (a) Bright-field microscopy image of a fully calcified *E. huxleyi* cell from a coculture. (b) Image of the same calcified cell (from a) under crossed polarizers. Coccoliths are evident covering the algal cell. Scale bar corresponds to 5 μm. (c) Bright-field microscopy image of a non-calcified (“naked”) *E. huxleyi* cell with attached bacteria in coculture. Arrow points to bacteria. (d) Image of the same cell (from c), under crossed polarizers showing no coccoliths. Scale bar corresponds to 5 μm. (e) Co-culture post algal death (left tube) and axenic algal culture (right tube), following centrifugation in a viscous solution that promotes coccolith separation from biomass (see Materials and Methods). Both samples exhibit a distinct white pellet indicative of coccoliths. (f) SEM image of the white pellet from the co-culture sample in (e). As can be seen, the pellet is indeed composed of coccoliths. Scale bar corresponds to 2 μm. (g) Comparison of the calcified cell portion in axenic algal cultures (“Alg”) and co-cultures (“Co”) at different time points. Both axenic cultures and co-cultures exhibit similar coccolith coverage at each time point. Data were collected from biological duplicates. Slight differences between samples were statistically insignificant as assessed by independent two-sample *t* test, assuming unequal variances with a significance level of α = 0.05. * Algal death in co-cultures occurred before day 15, therefore algal cells were not counted.

Next, we wished to examine whether bacteria influence algal coccolith coverage in the algal population. To this end, we quantified the number of naked and calcified algae in axenic versus co-cultures. Our results indicate that algal populations, both in axenic cultures and co-cultures, exhibit a similar proportion of calcified cells at each growth stage (Fig. 1g). Therefore, it appears that bacteria attach to a pre-existing sub-population of naked algal cells in co-cultures, while coccoliths production continues to occur in the adjacent calcified cells in the algal population. It remains to be determined whether naked algal cells do not produce coccoliths, or actually produce them but do not retain them on the cell surface.

Previously it was demonstrated that bacteria have a detectable impact on the physiology of coccolithophores and other micro-algae (Wang, Tomasch et al. 2014, Amin, Hmelo et al. 2015, Harvey, Deering et al. 2016, Segev, Wyche et al. 2016, Barak-Gavish, Frada et al. 2018, Whalen, Kirby et al. 2018). Importantly, algal alkenones that serve as a valuable proxy for sea surface temperature (SST) were altered in live algae due to the presence of bacteria (Segev, Castaneda et al. 2016). Given that algal calcification appears to proceed in co-cultures, the resulting coccoliths and their elemental composition can potentially be influenced by bacteria. To study whether bacteria affect coccolith elemental composition, we analyzed Sr/Ca ratios. Ratios of coccoliths Sr/Ca have been recognized as reliable and sensitive indicators of a range of temperatures and growth rates (Stoll and Schrag 2000, Stoll and Schrag 2001, Stoll, Klaas et al. 2002, Stoll, Rosenthal et al. 2002, Stoll, Ziveri et al. 2007, Saavedra-Pellitero, Baumann et al. 2017, Mejía, Paytan et al. 2018, Cavaleiro, Voelker et al. 2019). From a microbial perspective, growth rates of both algae and bacteria largely depend on temperature. Therefore, we first sought to explore the influence of different temperatures on dynamics in our algal-bacterial co-cultures. To this end, we cultivated cultures at different temperatures and found that the algal-bacterial interaction is highly affected by temperature (Fig. 2a, b, c). Higher temperatures resulted in faster growth rates (roughly 2.2 times faster at 22°C compared to 18°C, and 1.3 times faster at 18°C compared to 14°C. See Table 1), and in turn expedited microbial dynamics and resulted in earlier algal death (Fig. 2). Subsequent analyses of coccolith elemental composition revealed that the Sr/Ca ratio remained largely unaffected by the dynamic microbial interaction (*p* > 0.05). The Sr/Ca ratios in both axenic algal cultures and cocultures appear to correspond to temperature and growth rate regardless of microbial interactions (Fig. 3a). Elemental ratios of Mg/Ca, Mn/Ca and Ba/Ca were measured as well (Table S1, Figure S1), however yielded low reproducibility between biological replicates and exhibited no change across temperatures, treatments and time. Whether these results are due to technical or biological reasons is yet to be determined.

**Figure 2.**
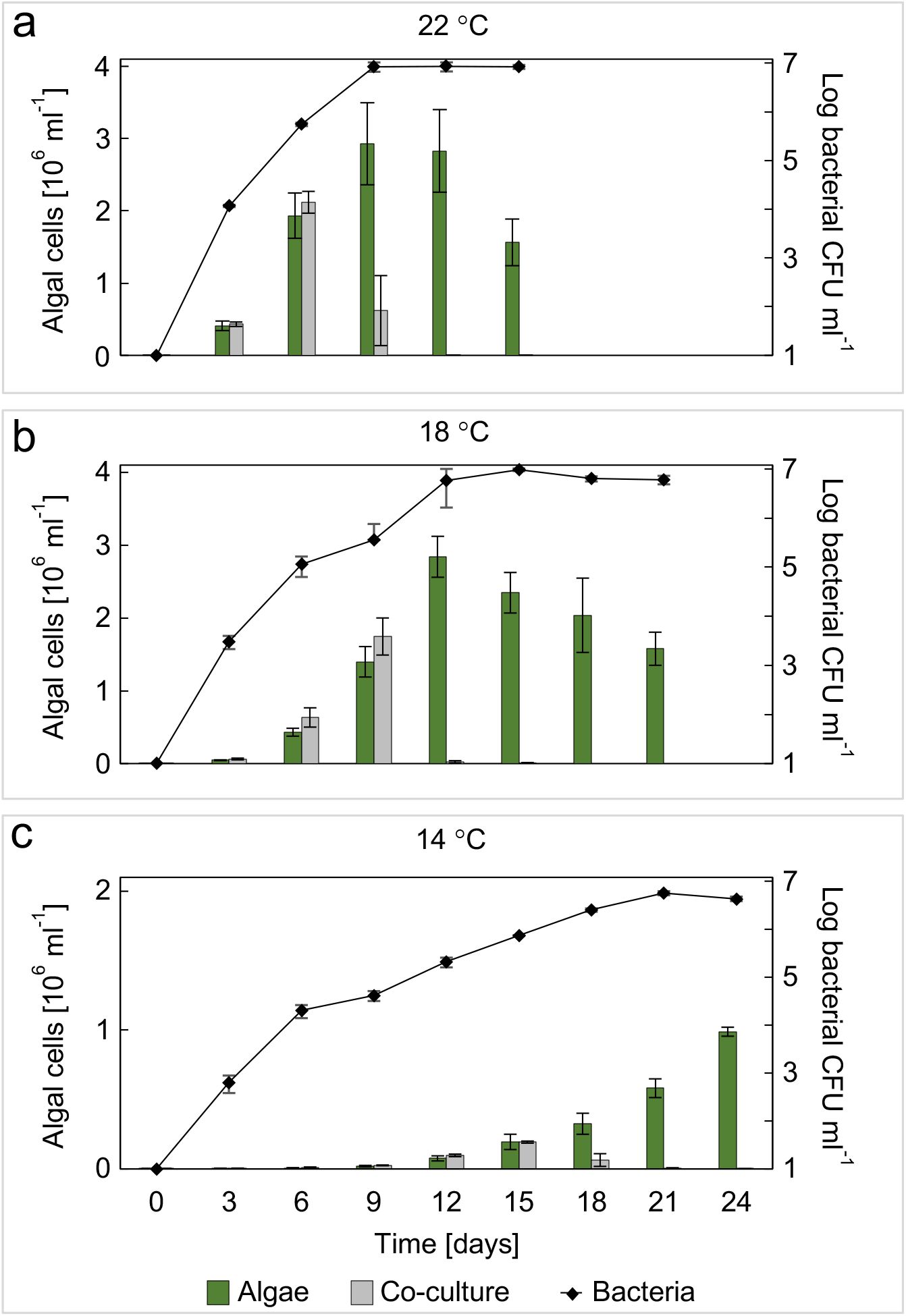
Rising temperatures expedite algal and bacterial growth dynamics. Growth curves of axenic algal cultures (green bars) and co-cultures (grey bars for algae, black line for bacteria). (a) Cultures were grown at 22°C and monitored over 15 days. (b) Cultures were grown at 18°C and monitored over 21 days. (c) Cultures were grown at 14°C and monitored over 24 days. Algal death in co-cultures occurred on day 10, 12 and 19 at 22°C, 18°C and 14°C, respectively. All data points represent the average value of three biological replicates. Error bars indicate the standard deviation between the three replicates.

**Table 1.** Algal growth rate per day at different temperatures. See materials and methods for calculation.

| Temperature [°C] | Growth rate [day<sup>-1</sup>] |
| --- | --- |
| 22 | 0.52 |
| 18 | 0.24 |
| 14 | 0.19 |

Growth rates and microbial dynamics vary considerably at different physiological stages of the cultures (Fig. 2). In the algal exponential phase, the growing algal population supports the growth of the bacterial population. As algae enter stationary phase, growth rate decreases and bacteria kill the algae (Segev, Wyche et al. 2016). Therefore, we next investigated whether microbial interactions have an effect on coccolith Sr/Ca at different time points of co-culturing. Coccoliths were harvested from two different time points during the cultivation of axenic and co-cultures. The chosen time points represent distinct physiologies during the algal-bacterial interaction; in the early time point (day 5 at 22°C and day 7 at 18°C) both algae and bacteria in co-cultures are exponentially growing. At the later time point, (day 11 at 22°C and day 14 at 18°C) bacteria kill their algal partners, while algae in the corresponding axenic cultures enter stationary phase. Of note, attempts to sample the first time point at 14°C were unsuccessful due to the very slow growth that resulted in insufficient coccolith yield, therefore only the later time point was sampled (day 20 at 14°C). Our analyses indicate that the presence of bacteria in co-cultures does not affect coccolith Sr/Ca ratio at different physiological stages (*p* > 0.05) (Fig 3b, c). Even though algal physiology is altered by bacteria during the algal-bacterial interaction (Fig. 2), the Sr/Ca ratio is unchanged. These observations reinforce the robustness of Sr/Ca ratio as an indicator of temperature and growth rate, independent of microbial interactions.

**Figure 3.**
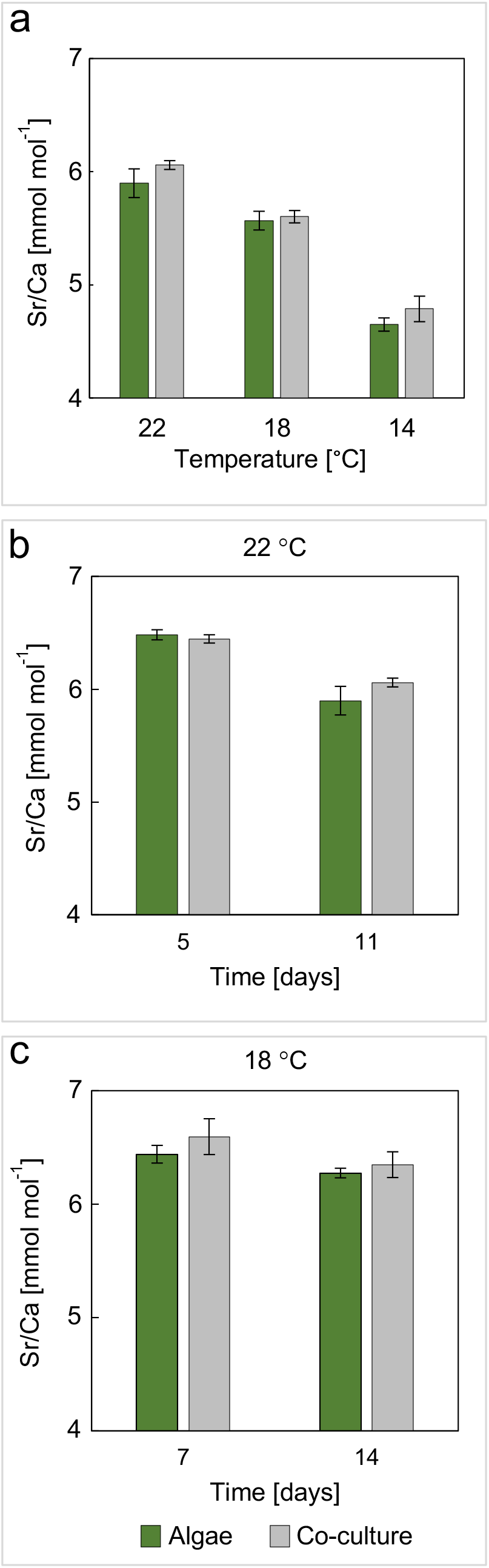
Coccolith Sr/Ca is influenced by growth rate and temperature, but not by microbial interactions. (a) Comparison of coccolith Sr/Ca between coccoliths from axenic algal cultures (green bars) and co-cultures (gray bars) at different temperatures indicates that higher temperatures result in higher Sr/Ca ratios. Measurements were conducted using cultures at their stationary growth phase (day 11 at 22°C, day 14 at 18°C and day 20 at 14°C). Differences between temperatures were statistically significant; *p* = 0.009 between 22°C and 18°C, *p* = 0.0005 between 18°C and 14°C (calculated according to axenic cultures values). No significant difference in Sr/Ca values was found between culture types at a given temperature. (b) Comparison of coccolith Sr/Ca at different stages of microbial growth at 22°C. Younger, exponentially-growing cultures (day 5) produced higher Sr/Ca ratios in comparison to older, stationary phase cultures (day 11). Differences between younger and older cultures were statistically significant with *p* = 0.001 (calculated according to axenic culture values). No significant difference in Sr/Ca ratios was observed between coccoliths from axenic algal cultures (green bars) and cocultures (gray bars). (c) Comparison of coccolith Sr/Ca at different stages of microbial growth at 18°C indicates no significant difference with *p* = 0.05 (calculated for axenic cultures).

## Discussion

The biological role of coccoliths is largely unknown (Müller 2019). Here we find that *E. huxleyi* strain CCMP3266 exhibits high heterogeneity in coccolith coverage, a phenotype inherent to this algal strain. The fact that bacterial attachment is restricted solely to naked algal cells, points to a possible and novel role of the coccosphere in algal-bacterial ecology.

Coccolithophores, similarly to other microorganisms, are part of a complex network of chemical interactions with other microbes, namely bacteria (Harvey, Deering et al. 2016, Segev, Wyche et al. 2016, Barak-Gavish, Frada et al. 2018, Whalen, Kirby et al. 2018). These interactions influence diverse aspects of algal physiology, resulting in changes in the composition and structure of different organic algal compounds such as alkenones, that are used as paleo-proxies (Segev, Castaneda et al. 2016, Fulton, Kendrick et al. 2017). Interestingly, viral infection of coccolithophores was also shown to influence algal alkenones during infection (Fulton, Kendrick et al. 2017). To the best of our knowledge, the current study is the first to explore coccolith elemental composition in the context of microbial interactions.

Previous studies demonstrated that algal-bacterial interactions influence algal alkenones (Segev, Castaneda et al. 2016), an established SST proxy (Brassell, Eglinton et al. 1986, Prahl and Wakeham 1987, Marlowe, Brassell et al. 1990). However, our results demonstrate no microbial influence on coccolith Sr/Ca ratio, another SST proxy. To understand the difference in the microbial influence on alkenones versus coccolith Sr/Ca, one should recognize the fundamental difference in the synthesis of the two proxies. Organic molecules, such as algal alkenones, are the product of regulated biosynthetic pathways that dictate the abundance and structure of the resulting compound (Rontani, Prahl et al. 2006, Zheng, Dillon et al. 2016). Therefore, a bacterial impact on algal physiology could have a general influence on cellular processes, including those related to alkenone biosynthesis. In contrast, coccolith CaCO_3_ crystals are precipitated directly from solution when conditions are favorable (Gal, Wirth et al. 2016). The final coccolith Sr/Ca ratio is thought to be controlled solely by thermodynamic and kinetic processes occurring inside the coccolith vesicle (Stoll, Rosenthal et al. 2002).

Similar to previous studies, we find a significant difference in the coccolith Sr/Ca ratio between cultures under different temperatures (Fig 3a). Previous studies with laboratory cultures and field observations have attributed variations in coccolith Sr/Ca to changes in growth rate and temperature (Stoll, Klaas et al. 2002, Stoll, Rosenthal et al. 2002, Stoll, Ziveri et al. 2007, Müller, Lebrato et al. 2014). However, it is still debated which of the two factors, growth rate or temperature, has a key control over coccolith Sr/Ca (Stoll, Klaas et al. 2002). In our experiments rising temperatures resulted in higher growth rates and elevated coccolith Sr/Ca ratio. It is therefore difficult to detangle, using our experimental system, the influence of each of the two variables on the final observed Sr/Ca.

Growth rates change significantly as cultures age. Laboratory cultures offer a unique opportunity to tightly monitor a synchronized microbial population throughout different growth stages. Interestingly, comparison of the exponential growth phase and the stationary phase in cultures cultivated at 22°C, revealed a measurable difference in coccolith Sr/Ca *(p* = 0.001). Previous studies described a correlation between coccolith Sr/Ca values and sea water Sr/Ca (Langer, Gussone et al. 2006). In our cultures, the Sr/Ca ratio of the growth medium at the end of culturing increases compared to the initial medium (Table S2), while the coccolith Sr/Ca decreases as growth progresses (Fig. 3b). Therefore, it appears that the observed differences in coccolith Sr/Ca between different growth phases are not the result of changing water chemistry.

Slight variability might be seen between results obtained from independent experiments (such as the data for 18°C in Fig. 3a compared with Fig. 3c). However, results obtained within a given experiment are highly reproducible (such as data shown at Fig. 3a at various temperatures). Of note, in our experimental system, coccoliths that are collected at each time point represent the cumulative sum of coccoliths that were produced during the cultivation period. It is tempting to hypothesize whether an even larger difference in coccolith Sr/Ca could be detected between growth stages by collecting only newly-produced coccoliths at each time point. Needless to say, that in natural microbial populations at sea, where algal communities are comprised of cells at different ages, such growth-phase-specific impacts are difficult to detect. Therefore, our experimental approach provides a good representation of natural coccolith-derived paleo-proxies, where different growth phases are presented collectively in sedimentary layers.

To conclude, it is becoming increasingly evident that coccolithophores have co-evolved with various microorganisms and that algal-bacterial interactions have been a driving force in the evolution of the interacting partners. The extensive fossil record produced by coccolithophores throughout the history of our planet, was produced while these algae were engaging in microbial interactions.

Our study highlights the key difference between organic and inorganic coccolithophore-derived paleo-proxies. While organic paleo-proxies are prone to biological influences during their production, inorganic paleo-proxies are mainly under thermodynamic and kinetic controls and are thus less sensitive to influences originating from microbial interactions.

## Acknowledgements

We are thankful to Jonathan Erez for insightful discussions and help during early stages of this research. We thank Itay Halevy and Assaf Gal for their valuable comments on the manuscript. We thank Tzachi Jacobson, Nir Galili, Ruth Yam and Aldo Shemesh for valuable technical advice and assistance. We thank Ophir Tirosh and Yigal Erel for their kind assistance with ICP-MS analyses. Finally, we are thankful to all members of the Segev lab for insightful discussions.

We gratefully acknowledge financial support from the Israeli Science Foundation (ISF 947/18), the Peter and Patricia Gruber Foundation, the Minerva Foundation with funding from the Federal German Ministry for Education and Research, and the Weizmann Sustainability and Energy Research Initiative (SAERI).

* Authors declare no conflict of interest.

## Materials and Methods

### Strains and culture conditions

The algal strain in this study was *E. huxleyi*CCMP3266 obtained from the National Center for Marine Algae and Microbiota, Maine, USA. Axenic algal cultures were inoculated with an initial cell concentration of 40 cells ml^-1^, and cultivated in 1 L glass Erlenmeyer flasks containing 0.5 L L1-Si growth medium prepared according to Segev et al., 2016 (1). L1-Si was prepared using 0.2 μm-filtered and autoclaved natural Mediterranean Sea water collected from Michmoret, Israel. Growth medium was filtered again prior to algal inoculation to remove particles that had formed during autoclavation. Flasks were cultivated in separate water baths maintained at 14, 18 and 22°C, under light/dark cycle of 16/8 hours with light intensity of 130 μm photons m^-2^s^-1^.

The bacterial species in this study was *P. inhibens* DSM17395, obtained from the DSMZ German Collection of Microorganisms and Cell Cultures, Braunschweig, Germany. To initiate bacterial cultures, frozen bacterial stocks were plated on 1/2 YTSS agar plates containing 2 gr yeast extract, 1.25 gr tryptone, 20 gr sea salts (Sigma Aldrich) and 16 gr agar (BD biosciences) in 1 L distilled water. Plates were incubated for 24 hours at 30°C. A single colony was transferred into liquid medium containing 10 ml of sea water supplemented with 5.5 mM glucose (Sigma Aldrich), 33 mM Na_2_SO_4_ (Merck), 5 mM NH_4_Cl (Sigma Aldrich) and 2 mM KH_2_PO_4_ (Carl Roth) (1). Liquid bacterial cultures were cultivated at 30°C for 24 hours, shaking at 130 rpm.

To initiate algal-bacterial co-cultures, after 4 days of cultivation algal cultures were inoculated with bacteria. Day 4 of algal growth, or upon addition of bacteria in co-cultures, is termed day 0 in both axenic cultures and co-cultures in all experiments. The bacterial inoculum was added at a final concentration of roughly 50 colony forming units (CFU) ml^-1^. Co-cultures were cultivated under the same conditions as their corresponding axenic algal cultures, as specified above.

Cultures intended for coccolith elemental analysis were grown in quadruplicates and were undisturbed until coccoliths were harvested. Cultures intended for algal and bacterial growth rate measurements and coccolith coverage assessment were homogenized by rigorous agitation every three days before a sample was taken for further analysis.

### Algal and bacterial cell enumeration

Algal cell density was measured using triplicate cultures on a Merck CellStream CS-100496 flow cytometer by plotting chlorophyll autofluorescence (excited at 561 nm, collected at 615-789 nm) against side scattering, and quantifying high chlorophyll events. For each sample, 50,000 events were recorded. Bacterial CFU in co-cultures was assessed by plating serial dilutions on 1/2 YTSS plates.

Doubling time of algal populations was calculated using the equation (4)

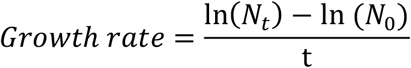

where *N*_0_ and *N_t_* denote the cell densities at the beginning and end of the logarithmic phase, and *t* is the phase length (expressed in days).

### Quantifying the portion of calcified algal cells

For assessing the portion of calcified cells in the algal population, 10 μl from duplicate cultures were loaded into a Bright-Line Hemacytometer and counted manually under a Nikon Eclipse Ni microscope equipped with Plan Fluor 40X lens. More than 300 cells were counted for every sample, except for day 3 in which only ~100 cells were counted due to low cell density. Cells were counted as calcified if they exhibited one or more attached coccoliths.

### Light microscopy

Light microscopy images were taken using Nikon Eclipse Ni microscope equipped with Plan Fluor 100X lens and polarizer. To reduce particle movement in the field of view, Liquid samples were loaded onto 1% agar pads prepared on a glass slide, and covered with a glass cover slip.

### Scanning electron microscopy

Scanning electron microscopy (SEM) images were taken using ZEISS™ Ultra 55 SEM, with SE2 detector and voltage of 5kV. Samples were coated with Iridium.

### Coccolith sample preparation

For coccolith purification, each culture was vigorously agitated to detach organic deposits that may have adhered to the glass vessel, and centrifuged into a single pellet consisting all of the organic and inorganic biomass that accumulated during growth. Each pellet was transferred into 2 ml Eppendorf tubes, and supernatant was discarded. Each pellet was resuspended in 1 ml of 100% Percoll (Sigma Aldrich) (2), and was homogenized at 30 Hz for 5 minutes using a RETSCH MM 400 mixer mill.

Homogenized pellets were centrifuged at 11000 RCF for 1 minute. The Percoll separates the organic fraction which floats at the top, from the coccolith fraction that remains at the bottom of the tube. The organic fraction and Percoll were discarded, and resuspension with Percoll was repeated. The coccolith fraction was washed twice using 18.2 MΩ ultrapure water (Merck Milli-Q IQ 7003) buffered to pH=9 using NH_4_OH.

Inspired by Blanco-Ameijeiras et al., 2012 (3) coccoliths were subjected to an oxidizing step for further removal of organic remains, with the following modifications; samples were treated with 1 ml 10% H_2_O_2_ (Fisher Scientific, trace analysis grade) buffered to pH=9 with NH_4_OH (Sigma Aldrich, trace metal basis), and heated to 80°C for 1 hour. The oxidizing procedure was conducted twice, followed by two washes with ultrapure water.

Next, coccoliths were treated for removal of possible metal oxides that could have deposited on the coccoliths surface. Following the protocol of Blanco-Ameijeiras et al., 2012 (3), samples were incubated in 1 ml 12% NH_2_OH·HCl (Sigma Aldric, 99% pure) buffered to pH=9 with NH_4_OH at room temperature for 24 hours. Samples were washed three times with ultrapure water, lyophilized and kept at room temperature until analyses.

### Inductively coupled plasma mass spectrometer (ICP-MS) measurements

Purified samples were weighed, dissolved in 2% HNO_3_ (Sigma Aldrich, trace metal basis) and diluted to final Ca^2+^ concentration of 50 ppm. Elemental analysis was conducted using Agilent 7500cx inductively coupled plasma mass spectrometer (ICP-MS) with an internal error typically below 5%. Each sample was measured three times and averaged. Prior to the analysis, the ICP-MS was calibrated with a series of multi-element standard solutions (Merck ME VI), and a series of Ca^2+^ standard solutions (SCP Science PlasmaCAL), both diluted in the same matrix as the samples.

To account for precision and possible drift during analysis, a series of control samples was examined at the beginning of the analysis, every 30 samples and at the end of the analysis. The series of control samples included a blank sample, standard reference samples (USGS SRS T-221, T-229), and several of the calibration standard solutions. To account for possible contamination during coccolith cleaning, several samples of CaCO_3_ powder (Sigma Aldrich, trace metal basis) were subjected to the entire coccolith cleaning process, and their elemental composition compared to uncleaned CaCO_3_. Prior to every experiment, all plastic equipment used for coccolith sample preparation and elemental analysis was submerged in 2% HNO_3_ for 48 hours and washed 5 times with ultrapure water. For monitoring elemental ratios and pH in the culture medium, measurements were conducted before culturing and at the end of culturing experiments (table S2).

### Statistics

Sr/Ca ratios between samples were regarded as different only if the difference was statistically significant, determined by independent two-sample *t* test, assuming unequal variances, with a significance level of α = 0.05.

