## Supplemental Figure S1 and Tables S1 and S2 for "Coccolith Sr/Ca is a Robust Temperature and Growth Rate Indicator that Withstands Dynamic Microbial Interactions"

Article title

**The following Supporting Information is available for this article:**

**Figure S1** Elemental ratios of coccolith $CaCO_3$.

**Table S1** Elemental ratios of coccolith $CaCO_3$.

**Table S2** Elemental composition and pH of culture media before and after microbial growth.

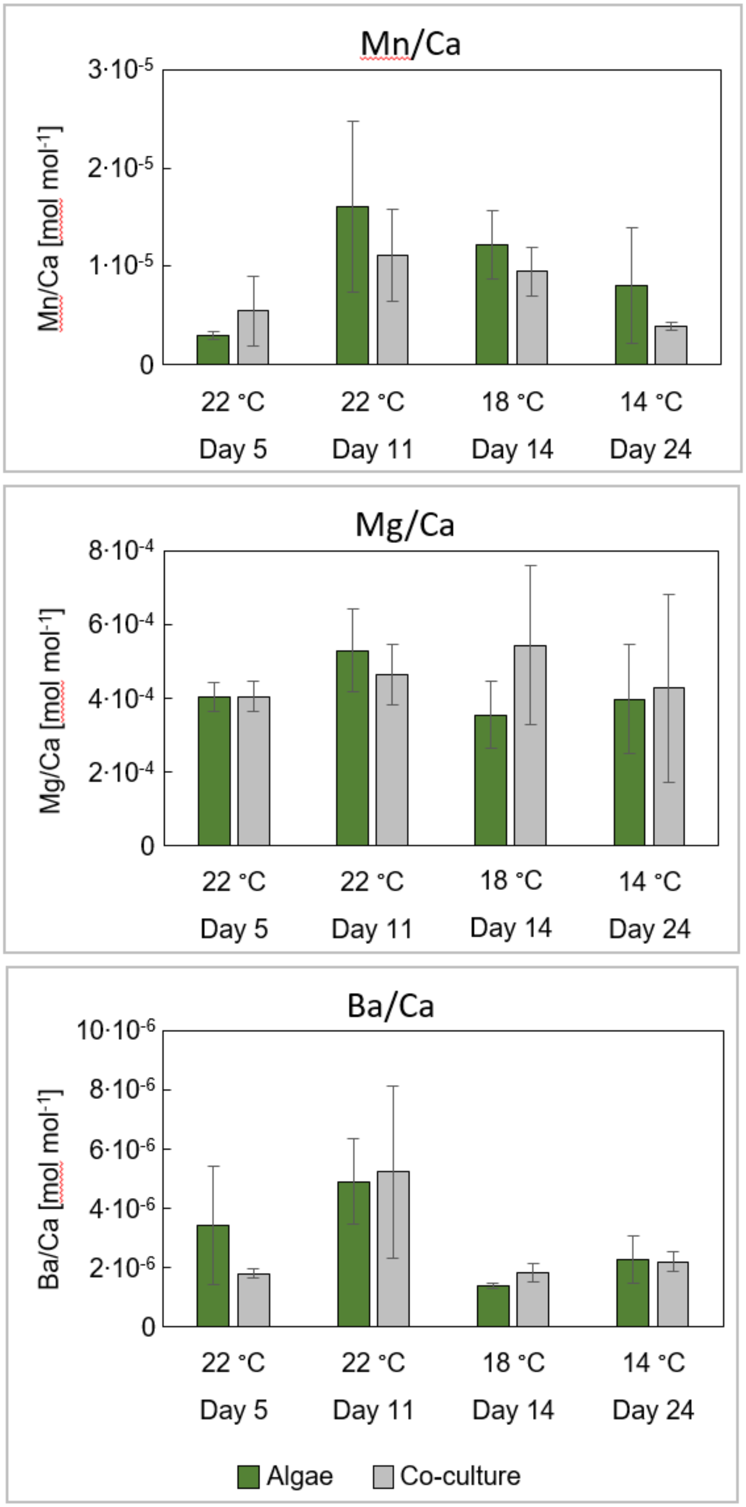


**Figure S1.** Elemental ratios of coccolith $CaCO_3$. Elemental ratios of coccolith Mg/Ca, Mn/Ca and Ba/Ca measured in parallel to the Sr/Ca values that are given in figure 3a, 3b.

**Table S1**. Elemental ratios of coccoliths $CaCO_3$

| Temperature [°C] | Culture type | Day | Mg/Ca | Mn/Ca | Sr/Ca | Ba/Ca |
|---|---|---|---|---|---|---|
| Measurements corresponding to samples presented in figure 3a, 3b, 4 | | | | | | |
| 22 | axenic | 5 | $4.01 \cdot 10^{-4}$ ($3.9 \cdot 10^{-5}$) | $2.95 \cdot 10^{-6}$ ($4.5 \cdot 10^{-7}$) | $6.48 \cdot 10^{-3}$ ($4.5 \cdot 10^{-5}$) | $3.43 \cdot 10^{-6}$ ($2.0 \cdot 10^{-6}$) |
| 22 | co-culture | 5 | $4.04 \cdot 10^{-4}$ ($4.0 \cdot 10^{-5}$) | $5.48 \cdot 10^{-6}$ ($3.5 \cdot 10^{-6}$) | $6.45 \cdot 10^{-3}$ ($3.7 \cdot 10^{-5}$) | $1.79 \cdot 10^{-6}$ ($1.7 \cdot 10^{-7}$) |
| 22 | axenic | 11 | $5.29 \cdot 10^{-4}$ ($1.1 \cdot 10^{-4}$) | $1.61 \cdot 10^{-5}$ ($8.8 \cdot 10^{-6}$) | $5.90 \cdot 10^{-3}$ ($1.3 \cdot 10^{-4}$) | $4.88 \cdot 10^{-6}$ ($1.4 \cdot 10^{-6}$) |
| 22 | co-culture | 11 | $4.63 \cdot 10^{-4}$ ($8.2 \cdot 10^{-5}$) | $1.11 \cdot 10^{-5}$ ($4.7 \cdot 10^{-6}$) | $6.06 \cdot 10^{-3}$ ($3.9 \cdot 10^{-5}$) | $5.21 \cdot 10^{-6}$ ($2.9 \cdot 10^{-6}$) |
| 18 | axenic | 14 | $3.54 \cdot 10^{-4}$ ($9.1 \cdot 10^{-5}$) | $1.21 \cdot 10^{-5}$ ($3.5 \cdot 10^{-6}$) | $5.57 \cdot 10^{-3}$ ($8.3 \cdot 10^{-5}$) | $1.39 \cdot 10^{-6}$ ($8.1 \cdot 10^{-8}$) |
| 18 | co-culture | 14 | $5.42 \cdot 10^{-4}$ ($2.1 \cdot 10^{-4}$) | $9.48 \cdot 10^{-6}$ ($2.5 \cdot 10^{-6}$) | $5.60 \cdot 10^{-3}$ ($5.5 \cdot 10^{-5}$) | $1.80 \cdot 10^{-6}$ ($3.2 \cdot 10^{-7}$) |
| 14 | axenic | 24 | $3.97 \cdot 10^{-4}$ ($1.5 \cdot 10^{-4}$) | $7.99 \cdot 10^{-6}$ ($5.9 \cdot 10^{-6}$) | $4.65 \cdot 10^{-3}$ ($5.8 \cdot 10^{-5}$) | $2.26 \cdot 10^{-6}$ ($7.9 \cdot 10^{-7}$) |
| 14 | co-culture | 24 | $4.27 \cdot 10^{-4}$ ($2.6 \cdot 10^{-4}$) | $3.87 \cdot 10^{-6}$ ($3.8 \cdot 10^{-7}$) | $4.79 \cdot 10^{-3}$ ($1.1 \cdot 10^{-4}$) | $2.18 \cdot 10^{-6}$ ($3.2 \cdot 10^{-7}$) |
| Measurements corresponding to samples presented in figure 3c | | | | | | |
| 18 | axenic | 7 | | | $6.44 \cdot 10^{-3}$ ($7.8 \cdot 10^{-5}$) | |
| 18 | co-culture | 7 | | | $6.59 \cdot 10^{-3}$ ($1.6 \cdot 10^{-4}$) | |
| 18 | axenic | 14 | | | $6.27 \cdot 10^{-3}$ ($4.2 \cdot 10^{-5}$) | |
| 18 | co-culture | 14 | | | $6.35 \cdot 10^{-3}$ ($1.1 \cdot 10^{-4}$) | |

Standard deviations (+/-) indicated in round brackets

**Table S2**. Elemental composition and pH of culture medium before and after algal growth

| sample type | pH | Na [ppm] | Mg [ppm] | Ca [ppm] | Mn [ppb] | Sr [ppm] | Ba [ppb] |
|---|---|---|---|---|---|---|---|
| initial medium | 8.30 | 11593 | 1333 | 113.10 | 59.20 | 9.70 | 19.51 |
| axenic | 8.48 (0.03) | 10844 (192) | 1271 (6.46) | 101.00 (1.66) | 31.30 (3.35) | 9.03 (0.14) | 18.71 (0.45) |
| co-culture | 7.95 (0.07) | 11123 (128) | 1283 (16.12) | 104.59 (3.58) | 47.87 (4.88) | 9.24 (0.19) | 18.70 (0.22) |

Values of axenic and co-cultures were collected in duplicates and averaged. Cultures were grown at 18 °C and sampled at the day of algal death in co-cultures. Standard deviations (+/-) indicated in round brackets.
